# Recognition of negative charge arrays by pleckstrin homology (PH) domains

**DOI:** 10.64898/2026.09.21.753330

**Authors:** Christopher K. Ng, James W. Murphy, Jeannine M. Mendrola, Michael Sarullo, Steven E. Stayrook, Kathryn M. Ferguson, Mark A. Lemmon

## Abstract

Pleckstrin homology (PH) domains are typically assumed to be phosphoinositide-binding modules, although most lack strong lipid specificity and their broader ligand repertoire remains poorly defined. We find that many yeast PH domains bind Nsr1p, the ortholog of nucleolin – also identified as a PH domain ligand – suggesting widespread recognition of negatively charged protein regions. The phosphatidylinositol 4,5-bisphosphate (PtdIns(4,5)*P*_2_)-binding PLCδ_1_PH domain also binds a highly phosphorylated region of IRBIT through the same site that recognizes PtdIns(4,5)*P*_2_. Using high-throughput integrated phosphopeptide (Hi-P) screening, we surveyed 38,624 mammalian sequences containing three documented phosphoserines.

PLCδ_1_-PH bound numerous phosphorylated and unphosphorylated acidic peptides, and phosphorylation generally strengthened pre-existing interactions rather than conferring strict specificity – without requiring a fixed phosphoserine spacing. Instead, favored phosphopeptides combined a key phosphoserine with upstream acidic residues. Structural modeling suggests that this phosphoserine occupies the canonical inositol phosphate binding pocket, while adjacent acidic residues make delocalized electrostatic contacts. Thus, PH domains can recognize various patterns of protein negative charge, expanding their potential regulatory roles beyond membrane targeting.

## INTRODUCTION

The pleckstrin homology (PH) domain is the 11^th^ most common domain in the human proteome,^1^ and among the most common signaling domains, with 250-270 examples across human proteins according to the SMART database.^2^ The first clues to PH domain function came from reports that certain examples bind phosphatidylinositol-4,5-bisphosphate (PtdIns(4,5)*P*_2_).^3,4^ In the case of the phospholipase-C-δ_1_ (PLCδ_1_) PH domain, the PtdIns(4,5)*P*_2_ headgroup is recognized stereospecifically.^5,6^ Subsequent studies then identified several PH domains that specifically recognize PtdIns(3,4,5)*P*_3_ and/or PtdIns(3,4)*P*_2_,^7,8^ linking a subgroup of PH domains to PI3-kinase signaling.^9^ Structural studies have shown how these PH domains engage inositol phosphate headgroups with specific spatial arrangements of phosphate groups.^10–14^ As a result of these initial findings, PH domains have been linked broadly to phosphoinositide binding.

Unlike Src homology 2 (SH2) and SH3 domains,^15^ which all appear to engage in binding to phosphotyrosine (SH2) and PxxP sequences (SH3),^16^ it soon became evident that the ability to specifically recognize phosphoinositides does not extend across the PH domain family.^17–19^ Rather, the PH domain appears to represent a conserved β sandwich protein scaffold or superfold^20^ that is particularly stable^21^ and can use different surfaces and loops to interact with multiple partners.^17,19,22,23^ Different PH domain-like modules have been shown to bind NPXpY sequences (the PTB or phosphotyrosine binding domain^24^), the small GTPase Ran (a PH-like domain in RanBP2^25^), or proline-rich peptides (the EVH1/WH1 domain^26^). Proteome-wide studies in *S. cerevisiae* further revealed that most PH domains from yeast do not bind strongly or selectively to phosphoinositides.^18^ There have been numerous reports of PH domain binding to acidic protein sequences. The IRS-1 and IRS-2 PH domains were found in yeast 2-hybrid studies to bind glutamate/aspartate rich regions from nucleolin, lon protease, and other proteins.^27^ Similarly, the PH domain from the TFIIH p62/Tfb1 subunit binds E/D rich peptides from the transactivation domains of p53 or TFIIEα.^28^ Interestingly, the ceramide transfer protein (CERT) PH domain binds intramolecularly to a phosphorylated serine-repeat motif, blocking its ability to interact in trans with phosphoinositide-containing membranes^29^ – echoing early suggestions by Saraste and colleagues that PH domains bind phosphorylated proteins.^30^

PH domains that specifically recognize phosphoinositides are strongly electrostatically polarized^12^ and possess a conserved pattern of basic residues (**KX_n_[K/R]XR**) centered on their β1/β2 loop^31–34^ that engage the inositol phosphate as shown in Figure 1A. Other PH domains appear to retain differently arranged sets of basic residues and bind PtdIns(4,5)*P*_2_ at a different site (as seen for β-spectrin^35^), and most appear to retain the property of electrostatic sidedness.^18,20^ Studies with a different inositol phosphate-binding protein, the Ins(1,4,5)*P*_3_ receptor identified a binding partner that may bridge these ligand classes. IRBIT (for IP₃R-binding protein released with Ins(1,4,5)*P*_3_), directly competes with Ins(1,4,5)*P*_3_ for binding to the endoplasmic reticulum Ins(1,4,5)*P*_3_ receptor,^36^ through a multiply phosphorylated region in the IRBIT amino-terminal region thought to mimic Ins(1,4,5)*P*_3_.^37^ Here, we asked whether ligands for PH domains may have similar phosphopeptide/inositol phosphate ‘duality’, by asking whether PH domains that do-and do not appear to specifically recognize phosphoinositides can also bind multiply phosphorylated peptides. We screened the ability of different PH domains – including the PtdIns(4,5)*P*_2_-binding PLCδ_1_ PH domain – to bind different protein sequences. We show that acidic phosphopeptide binding is at least as common a PH domain property as phosphoinositide binding, and has many structural parallel, combining stereospecific charge recognition and delocalized electrostatic attraction. These findings have important implications for the broader functions of PH domains, and suggest approaches for investigating understudied PH domain that currently have no known function.

**FIGURE 1.**
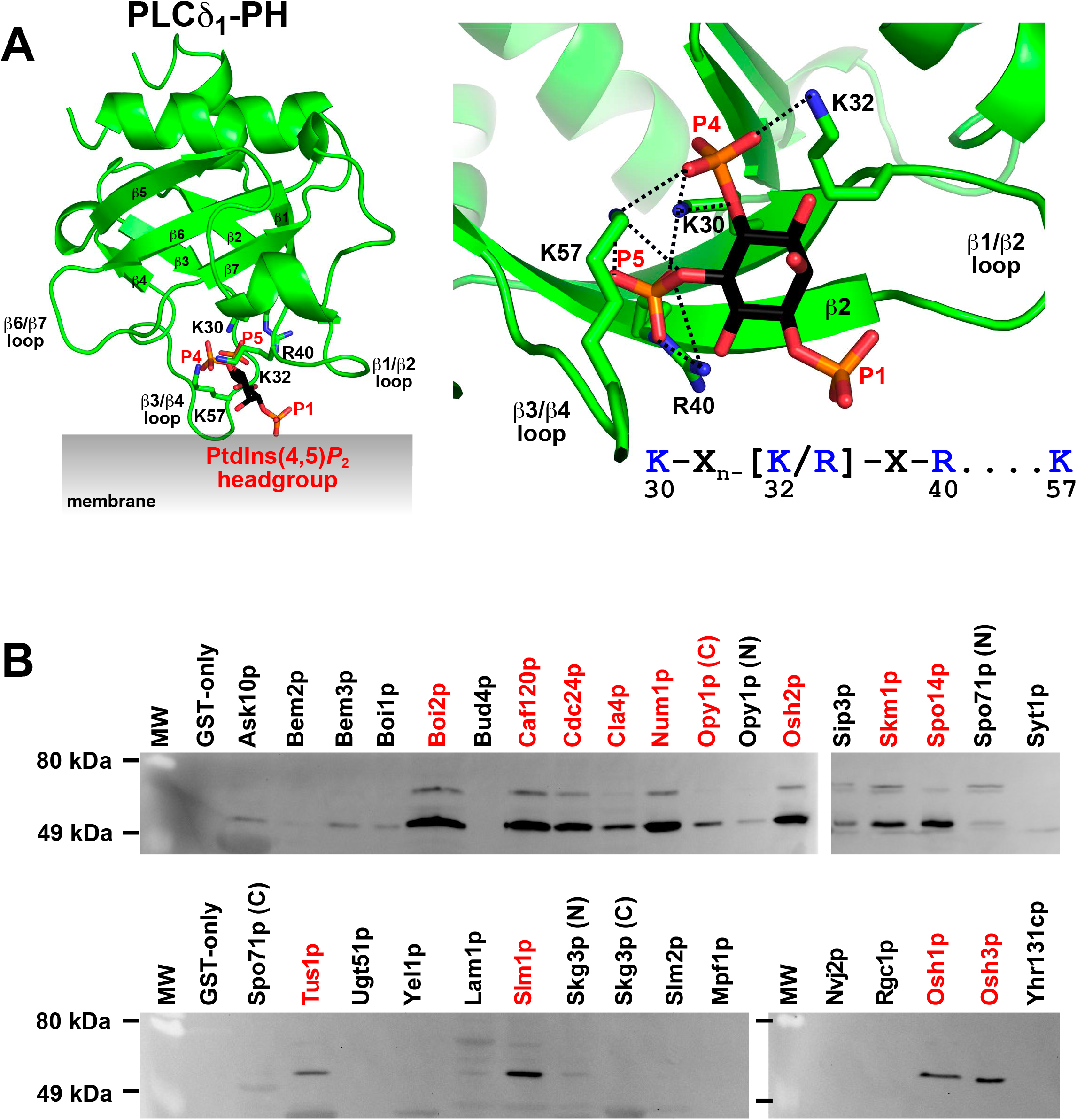
PH Domain recognition of PtdIns(4,5)*P*_2_ headgroup and Nsr1p **A.** Structure of PLCδ_1_-PH bound to Ins(1,4,5)*P*_3_, the headgroup of PtdIns(4,5)*P*_2_. *Left*, the PH domain β-sandwich is shown in green above a schematic membrane, engaging Ins(1,4,5)*P*_3_ (black, orange and red) placed as the PtdIns(4,5)*P*_2_ headgroup (shown as sticks). Strands β1 through β7 in the sandwich are marked, and basic residues in the PH domain β1/β2 and β3/β4 loops that engage Ins(1,4,5)*P*_3_ phosphates are shown. From PDB ID: 1MAI.^12^ *Right*, a close-up view of Ins(1,4,5)*P*_3_ bound between the β1/β2 and β3/β4 loops of PLCδ_1_-PH (PDB ID: 1MAI^12^), ‘clamped’ in by the basic side chains of K30, K32 and R40 in the β1/β2 loop and K57 in the β3/β4 loop shown in sticks. Dashed lines indicate polar contacts. The Ins(1,4,5)*P*_3_ phosphate groups are labeled P1, P4, and P5. The basic residue pattern is shown below, with the corresponding PLCδ_1_ residue numbers. **B.** Pulldown of TAP-tagged Nsr1p from yeast lysate by GST fusions of the indicated yeast PH domains immobilized on glutathione-Sepharose beads. Equivalent amounts of lysate were incubated with each bait, and recovered proteins were analyzed by SDS-PAGE and immunoblotting with a TAP antibody. GST-only beads served as a negative control. Red labels identify the 13 PH domains that precipitated Nsr1p among the 33 tested. N and C distinguish the N-terminal and C-terminal PH domains of proteins containing two domains. MW, molecular-weight markers.

## RESULTS AND DISCUSSION

### Many PH domains bind a nucleolin orthologue

Reports that multiple PH domains can bind robustly to proteins with highly acidic stretches – such as IRS-1/2 PH domains binding to nucleolin^27^ – led us to assess the ability of all *S. cerevisiae* PH domains^18^ to bind the yeast equivalent of nucleolin, Nsr1p (YGR159C).^38^ As shown in Figure 1B, at least 13 of the 33 GST-fused *S. cerevisiae* PH domains^18^ were able to precipitate Nsr1p with a tandem affinity purification (TAP) tag^39^ from a yeast cell lysate. This result suggested that Nsr1p binding is actually a more common property of yeast PH domains that phosphoinositide binding. Unexpectedly, the set of PH domains that bind Nsr1p in this assay included most of those that we previously found to bind phosphoinositides. In older studies, we had also identified the highly phosphorylated NOLC1/NOPP140 protein^40^ as a robust binding partner for the RAS-GAP/RASA1^41^ PH domain (which also binds phosphoinositides^42^) based on bacterial expression cloning. Collectively, these findings suggest a preponderance of cases in which both phosphoinositides and negatively charged regions of proteins bind to certain PH domains. In some cases, these two types of ligand may compete to regulate their host proteins, as suggested for CERT^29^ and AKT,^43^ for example.

### PLCδ_1_ PH domain binds IRBIT

Studies of IRBIT binding to the Ins(1,4,5)*P*_3_ receptor^36,37^ showed that it competes with the inositol phosphate ligand for its binding site on the Ins(1,4,5)*P*_3_ receptor. IRBIT is an S-adenosylhomocysteine hydrolase homolog with an amino-terminal extension that contains a multiply phosphorylated serine-rich region that is thought to mimic Ins(1,4,5)*P*_3_ to as a “pseudoligand”.^36,37^ Indeed, the same mutations in the Ins(1,4,5)*P*_3_ receptor impair binding of both phosphorylated IRBIT and Ins(1,4,5)*P*_3_.^36^ Since the PLCδ_1_ PH domain binds Ins(1,4,5)*P*_3_ with high affinity (*K*_d_ = 210 nM^6^) and stereospecificity,^6,12^ we asked whether it also binds IRBIT.

As shown in Figure 2A, IRBIT produced in Sf9 cells binds robustly to a GST-fusion of the PLCδ_1_ PH domain in surface plasmon resonance (SPR) studies, with an apparent *K*_d_ value in the 0.33 μM range. Binding is substantially diminished when the seven serines noted in the sequence at the top of Figure 2A (serines 66, 68, 70, 71, 74, 76, and 77) are replaced with alanines (IRBIT-A_7_), but is largely retained when they are replaced instead with glutamates (IRBIT-E_7_). Similarly, a GST/PLCδ_1_-PH fusion protein robustly pulled down histidine-tagged wild-type IRBIT overexpressed in Sf9 cells, but not in *E. coli*, mirroring the report of Ando *et al*.^36^ for IRBIT binding to the Ins(1,4,5)*P*_3_ receptor – showing dependence of PH domain binding by IRBIT on its phosphorylation. These experiments also revealed that the yeast Boi2p PH domain (from YER114C), which binds phosphoinositides robustly, also binds IRBIT in a phosphorylation-dependent manner.^18^ By contrast, the PH domain from dynamin-1, which binds only very weakly to phosphoinositides,^44,45^ fails to interact robustly with IRBIT whether produced in Sf9 cells or *E. coli*.

**Figure 2.**
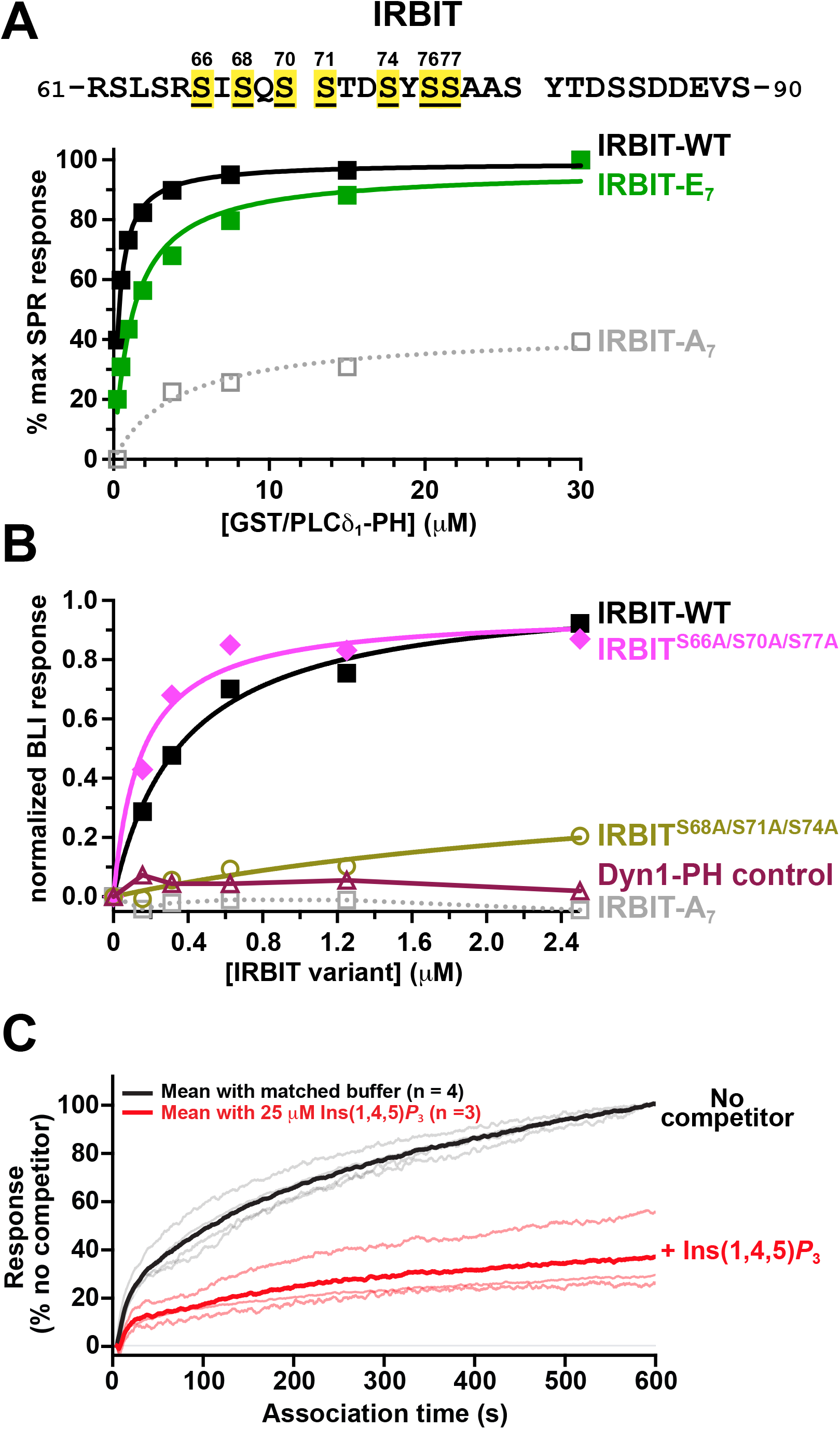
IRBIT binding to PLCδ_1_-PH depends on its serine-rich region **A.** Surface plasmon resonance (SPR) measurements of GST/PLCδ_1_-PH binding to immobilized full-length His_6_-tagged IRBIT produced in Sf9 cells. Wild-type IRBIT (IRBIT-WT; black) is compared with variants in which the seven serines highlighted in yellow and numbered in the IRBIT N-terminal sequence fragment shown at the top of the figure were replaced by alanines (IRBIT-A_7_; gray) or glutamates (IRBIT-E_7_; green). Failure to phosphorylate these serines also prevents binding. Responses were corrected by subtraction of a mock-coupled surface and expressed as a percentage of the maximal SPR response. These experiments are representative of at least three repeats. **B.** Biolayer interferometry (BLI) concentration/response measurements for soluble full-length IRBIT variants binding to GST/PLCδ_1_-PH immobilized on Octet anti-GST biosensors. IRBIT-WT (black), S66A/S70A/S77A-mutated IRBIT (magenta), S68A/S71A/S74A-mutated IRBIT (olive), and IRBIT-A_7_ (gray) are compared. Immobilized GST/Dyn1-PH (maroon) provides a low-binding control. Individual points show the mean response during the final 20 s of association in the BLI experiment, after subtraction of the corresponding response for equivalent sensors on which GST-only had been immobilized. Responses were normalized to the mean fitted B_max_ for IRBIT-WT and IRBIT^S66A/S70A/S77A^ in the same experiment. The data shown are representative of three independent experiments. **C.** BLI association traces for binding of IRBIT-WT from a 5 μM solution to sensor-immobilized GST/PLCδ_1_-PH with no soluble competitor (in matched buffer; black/gray) or in the presence of 25 μM Ins(1,4,5)*P*_3_ (red) as competitor. Traces were aligned at association onset, corrected by subtraction of signal from a matched GST-only sensor (and competitor-only signals), and normalized to the endpoint obtained with no competitor Fainter lines represent individual experiments; bold lines show means (no competitor, n = 4; Ins(1,4,5)*P*_3_, n = 3). One sensor was used per condition within each experiment.

To investigate IRBIT binding to the PLCδ_1_ PH domain in more detail, we immobilized a wild-type GST/PLCδ_1_-PH fusion protein on sensors for biolayer interferometry (BLI; see STAR Methods) and tested its ability to bind different IRBIT variants produced in Sf9 cells (Figure 2B). We observed robust binding of wild-type IRBIT to the PLCδ_1_ PH domain using BLI (apparent *K*_d_ = 0.42±0.22 μM), but not to the negative control dynamin-1 PH domain or to IRBIT-A_7_. Ando *et al*.^36^ reported previously that mutating S68, S71 or S74 in the IRBIT amino-terminal region sequence – **_62_-SLSRSISQS STDSYSSAAS YTDSSDDEVS-_90_**– to alanine or glycine abolished binding to the Ins(1,4,5)*P*_3_ receptor. We found that mutating these three residues to alanine also abolished IRBIT binding to PLCδ1-PH (Figure 2B). By contrast, leaving these three serines intact (and phosphorylatable), but replacing serines 66, 70, and 77 with alanines kept PLCδ_1_-PH binding intact (apparent *K*_d_ = 1.03±0.77 μM). Any triple serine-to-alanine mutation that included S68, S71 and/or S74 also abolished IRBIT binding to the PLCδ_1_ PH domain. Importantly, we found that adding soluble Ins(1,4,5)*P*_3_ as a competitor could prevent IRBIT binding to immobilized GST/PLCδ_1_-PH (Figure 2C), arguing that – as with the Ins(1,4,5)*P*_3_ receptor,^36^ phosphorylated IRBIT competes with Ins(1,4,5)*P*_3_ for binding to the PLCδ_1_ PH domain.

### Bacterial assay for PLCδ_1_-PH/IRBIT interaction

Based on the above results, we reasoned that the multiply phosphorylated serine-rich amino-terminal region of IRBIT may mimic Ins(1,4,5)*P*_3_ to bind strongly to PLCδ_1_-PH as previously suggested for the Ins(1,4,5)*P*_3_ receptor.^46^ With a view to screening for serine-phosphorylated peptides capable of mimicking the stereochemistry of Ins(1,4,5)*P*_3_, we turned to an *E. coli*-based High-throughput integrated Phosphopeptide screening (Hi-P) approach described by Barber *et al*.^47^ Hi-P uses a single-plasmid library encoding many mammalian phosphopeptides to screen for serine-phosphorylation-dependent protein interactions based on bimolecular fluorescence complementation (BiFC) arising from (re)association of two fragments of a split mCherry protein.^48^ We implemented the Hi-P platform in the genomically recoded *E. coli* strain C321.ΔA.^49^ Initial optimization experiments directed us to use GST/PLCδ_1_-PH fused to the N terminus of a C-terminal mCherry fragment (mCherryC) and peptide fused to the C-terminus of the N-terminal mCherry fragment, both in the bicistronic reporter plasmid. UAG codons were incorporated in the peptide at the desired phosphoserine locations. When expressed with the pSer orthogonal translation system (SepOTSλ), phosphoserines (pSer) are incorporated at these positions in the pSer arm of the experiment, whereas co-expression of the amber suppressor tRNA^Ser^*_CUA_* (*supD* tRNA) in the serine control (nSer) arm of the screen incorporates unmodified serine at these sites.^50,51^

We first asked whether this platform could be used to detect interaction of GST/PLCδ_1_-PH-mCherryC with a phosphorylated IRBIT peptide fused to mCherryN. We used an IRBIT peptide (IR20) spanning residues 51-91 of human IRBIT, which has the sequence (when phosphorylated): **`FTKFPTKTGRRSLSRSI(pSer)QS(pSer)TD(pSer)YSSAASYTDSSDDEVSP`**. UAG codons at positions corresponding to serines 68, 71, and 74 direct pSer incorporation at those positions. As shown in Figure 3A, plate reader-measured OD_620_-normalized mCherry fluorescence for cultures expressing GST/PLCδ1-PH-mCherryC was substantially (4-5 fold) higher with the phosphorylated than non-phosphorylated IRBIT peptide fused to mCherryN. Fluorescence signals (and phosphorylation dependence) were similar to those obtained in parallel positive control experiments using the 14-3-3β/phospho-palladin interaction described by Barber et al. identified using Hi-P^47^ Another phosphoinositide-binding PH domain from *S. cerevisiae* Osh2p^18^ also showed evidence for phosphorylation-dependent interaction with the IR20 peptide in this assay, consistent with the hypothesized ability of pSer-containing IR20 to mimic inositol phosphates. Importantly, phosphorylation-dependent fluorescence recovery with the singly phosphorylated palladin peptide was much less extensive with the PLCδ_1_ or Osh2p PH domains than for 14-3-3β, consistent with a requirement for multiple phosphoserines in PH domain binding to IR20 and IRBIT.

**Figure 3.**
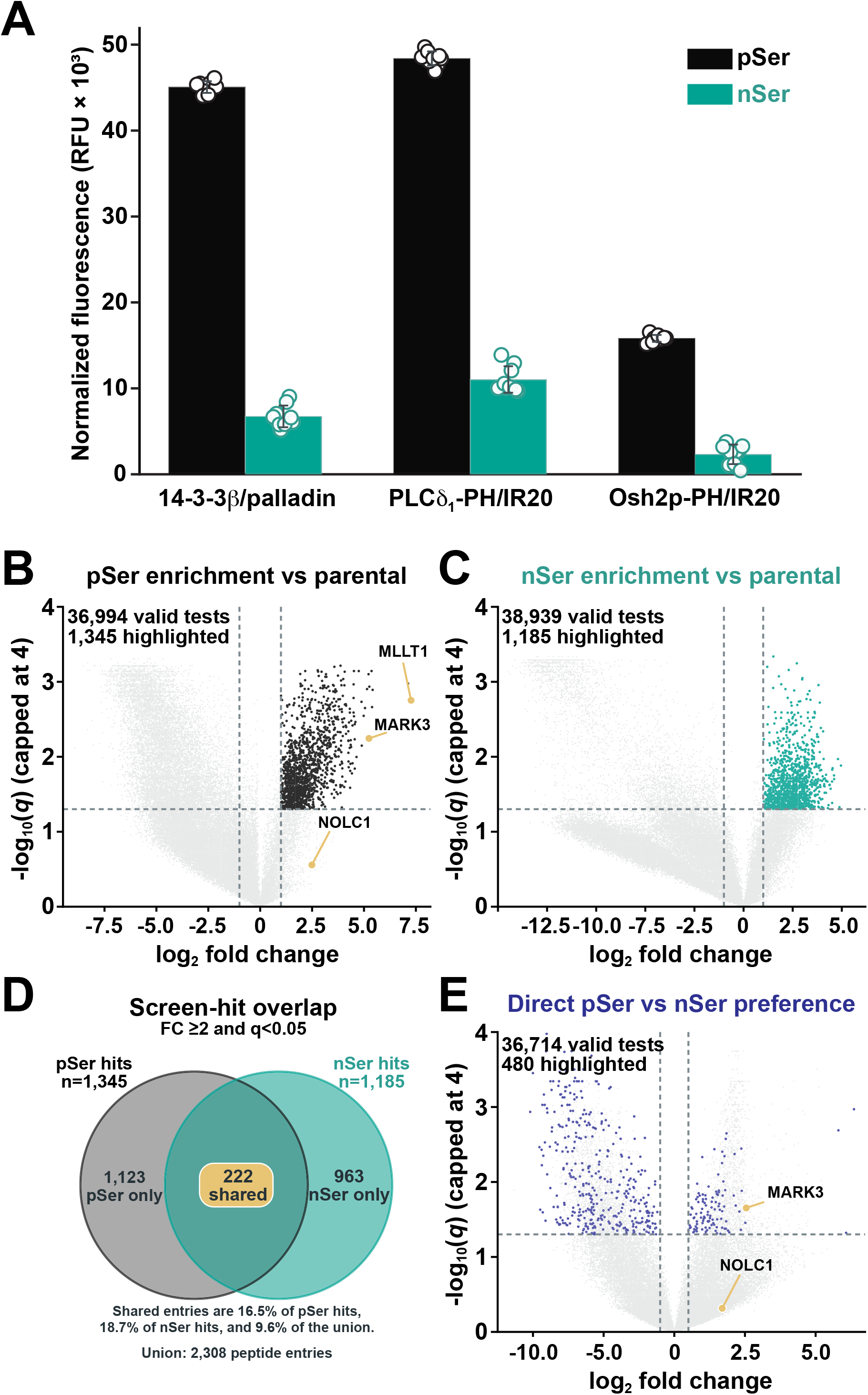
Hi-P analysis of peptide binding to PLCδ1-PH **A.** Representative split-mCherry reporter responses for the 14-3-3β/palladin benchmark from Barber et al.^47^ and GST-tagged PLCδ_1_-PH or Osh2p-PH paired with the IRBIT-derived IR20 peptide described in the text. Programmed UAG sites encode phosphoserine in the pSer condition (SepOTSλ; black) or serine in the nSer condition (*supD*; teal). IR20 spans IRBIT residues 51–91, with programmed UAG sites at S68, S71, and S74. Bars show means, whiskers show standard deviations (SD), and open circles show technical wells (n = 9 per condition). Fluorescence values were corrected for LB-only background and normalized by the corresponding blank-corrected OD_620_ value. (**B** and **C**) Volcano plots of peptides enriched relative to their matched unsorted parental populations in pSer (**B**) and nSer (**C**) enrichment screens. Each point represents an annotated library entry. Black (**B**) and teal (**C**) points satisfy fold enrichment (FC) ≥2, Benjamini-Hochberg adjusted q < 0.05, and non zero selected signal, which identify 1,345 pSer and 1,185 nSer hits. Gold points in **B** highlight peptide entries discussed in the text. **D.** Overlap of the within-screen hit sets, with 222 shared entries among 2,308 entries identified in either screen. **E.** Direct comparison of enrichment in the pSer and nSer screens. The horizontal axis shows log₂(FCpSer/FCnSer). Blue points satisfy a ≥2-fold difference, direct q < 0.05, and the within-screen hit criteria in the favored arm. The 480 highlighted entries comprise 108 pSer-preferred and 372 nSer-preferred entries. Gold points highlight entries discussed in the text For **B** to **E**, each arm comprised four independent biological replicates, with enrichment measured after two sorting rounds relative to the matched parent. Tests required at least three informative selected-parental pairs per arm. Within-screen comparisons used two-sided one-sample t tests of replicate log2 enrichment against zero. Direct comparisons used two-sided Welch t tests. Benjamini-Hochberg correction was applied separately to the three test families. In **B**, **C**, and **E**, valid-test counts are indicated and dashed lines mark the twofold and q = 0.05 thresholds.

### Screen for PLCδ_1_-PH-binding phosphopeptides

Having demonstrated that phosphorylation-dependent association of PLCδ_1_-PH with IR20 can be detected using Hi-P, we next designed a screen to investigate preferred phosphosite spacing, local sequence context and charge distribution for this interaction. Rather than screen randomized sequences in efforts to establish a three-phosphoserine motif for PLCδ_1_-PH that may reflect unnatural sequences, we treated the **pS_68_xxpS_71_xxpS_74_** pattern in IRBIT implicated by Ando *et al*.^36^ and in Figure 2C as a starting hypothesis. We then designed a library to ask whether PLCδ_1_-PH binds selectively to peptides represented in existing mammalian databases that contain three documented phosphoserines with specific flanking residues, intervening residues, spacing, or other sequence features. We filtered documented human, mouse, and rat phosphoserine sites from PhosphoSitePlus^52^ into 11 pre-specified classes based on phosphoserine spacing. With **J** denoting a site programmed to be occupied by pSer or Ser, 9 classes of sequence contained every **J***x***J***y***J** combination in which *x* and *y* were 0, 1, or 2 intervening residues (IR20 has a **J**2**J**2**J** pattern). **J**3**J**3**J** and **J**4**J**4**J** sequences provided two additional symmetric classes. For each such motif identified in the databases, sequence windows of ±15 residues and ±20 residues either side of the central phosphoserine (**J**) were generated when sequence context permitted. The resulting 31-41 amino acid library entries were codon optimized and UAG codons were included at the pSer/Ser sites. The final annotation table contained 41,848 phosphorylated sequences in PhosphoSitePlus, represented after removal of protein sequence redundancy by 38,624 unique 109-184 bp oligonucleotide designs (including linkers). All sequences were synthesized in duplicate commercially (by Agilent), pooled, and ligated into the bicistronic Hi-P vector fused to the mCherryN carboxy-terminus. The resulting bicistronic vectors thus each contained GST/PLCδ_1_-PH-mCherryC plus one library member fused to mCherryN. After construction (see Methods), the library was introduced in separate transformations into C321.ΔA cells carrying SepOTSλ or *supD*. Four unsorted transformed cell populations showed non-zero representation of 37,635, 37,627, 37,574, and 37,705 unique sequences, corresponding to 97.3% to 97.6% of the 38,624 unique DNA sequences. Starting abundances of individual library members varied for each independent population, so every selected population was analyzed relative to its own unsorted parent in the experiments below. Importantly, this approach bases the screened library on actual mammalian sequences with known phosphorylation sites, rather than abstract random sequences that may have no biological corollary.

We screened both pSer (with SepOTSλ) and nSer (with *supD*) in four independent biological replicates each. In each case, following overnight reporter induction (see Methods), the top 2% of mCherry events were collected by fluorescence activated cell sorting (FACS) with a target of 100,000 events. Recovered cells from the top 2% were expanded, re-induced, and subjected to a second top 2% FACS sort. DNA extracted from the doubly selected populations and their matched unsorted parent pools was then subjected to next generation sequencing. Sequences that were selected for or against in the sorted population compared with unsorted cell population were identified for each screen replicate, with statistical analysis restricted to those sequences for which enrichment/de-enrichment could be determined in at least three biological repeats for the pSer or nSer screen. The statistical criteria outlined in Methods allowed us to determine direct p values in three separately corrected Benjamini–Hochberg families for 36,994 pSer protein sequences, 38,939 nSer sequences, and 36,714 direct pSer versus nSer comparisons (Figure 3B-E). Defining a positive within-screen hit as requiring at least twofold enrichment, q<0.05, and non-zero selected signal in at least one replicate identifies 1,345 pSer hits (Figure 3B) and 1,185 nSer hits (Figure 3C), corresponding to 1,282 and 1,143 unique translated peptides, respectively. The pSer hits map to 278 gene labels, whereas the nSer hits map to 475. Of the 1,345 hits in the pSer screen shown in Figure 3B, 1,123 are unique to the pSer screen, and 222 are shared with the nSer screen (Figure 3D).

We also compared replicate-level enrichment vectors from the independent pSer and nSer series using a two-sided Welch test (Figure 3E). Among 36,714 eligible direct tests, 4,362 entries differed by at least twofold with direct q<0.05: 1,627 in the pSer-preferred direction and 2,735 in the nSer-preferred direction. Many represented differential depletion or weak enrichment, so direct preference was anchored in most cases to a positive hit within the pSer or nSer screen. Importantly, across the two screens, strict pSer selectivity was essentially never seen, with pSer-favored entries retaining measurable nSer signal in every case. This result indicates that pSer incorporation typically strengthened an existing response rather than eliciting new interactions.

### Features of PLCδ_1_-PH binding pSer and nSer peptides

Contrary to our initial hypothesis, the Hi-P results do not support the idea that serine-phosphorylated peptides mimic the stereochemistry of Ins(1,4,5)*P*_3_ to bind the PLCδ_1_ PH domain. One clear piece of evidence for this is the fact that all 11 of the pSer spacings in the initial library were represented both among pSer hits (Figure 4A) and among pSer-‘preferred’ entries for which the pSer/nSer fold change is ≥2 with q<0.05. After exact translated peptide duplicates were removed from each set, the compact **J**0**J**0**J** class that has three consecutive programmed pSer sites was slightly enriched compared to its representation in the parental library among pSer hits (by 1.4-fold), nSer hits (by 1.5-fold) and pSer-preferred sequences (by 2.8-fold). By contrast, the IRBIT-associated **J**2**J**2**J** class was not significantly enriched among any set of hits. Although certain spacings were under-represented across the relatively small number (108) of pSer-preferred sequences, it is clear from Figure 4A that no phosphoserine spacing class defines an obligatory geometry. These data therefore suggest that the peptide sequence requirement for binding to PLCδ_1_-PH in the Hi-P assay are substantially less stringent than suggested by the inositol phosphate mimicry or Ins(1,4,5)P3 ‘pseudoligand’^36,46^ hypothesis.

**Figure 4.**
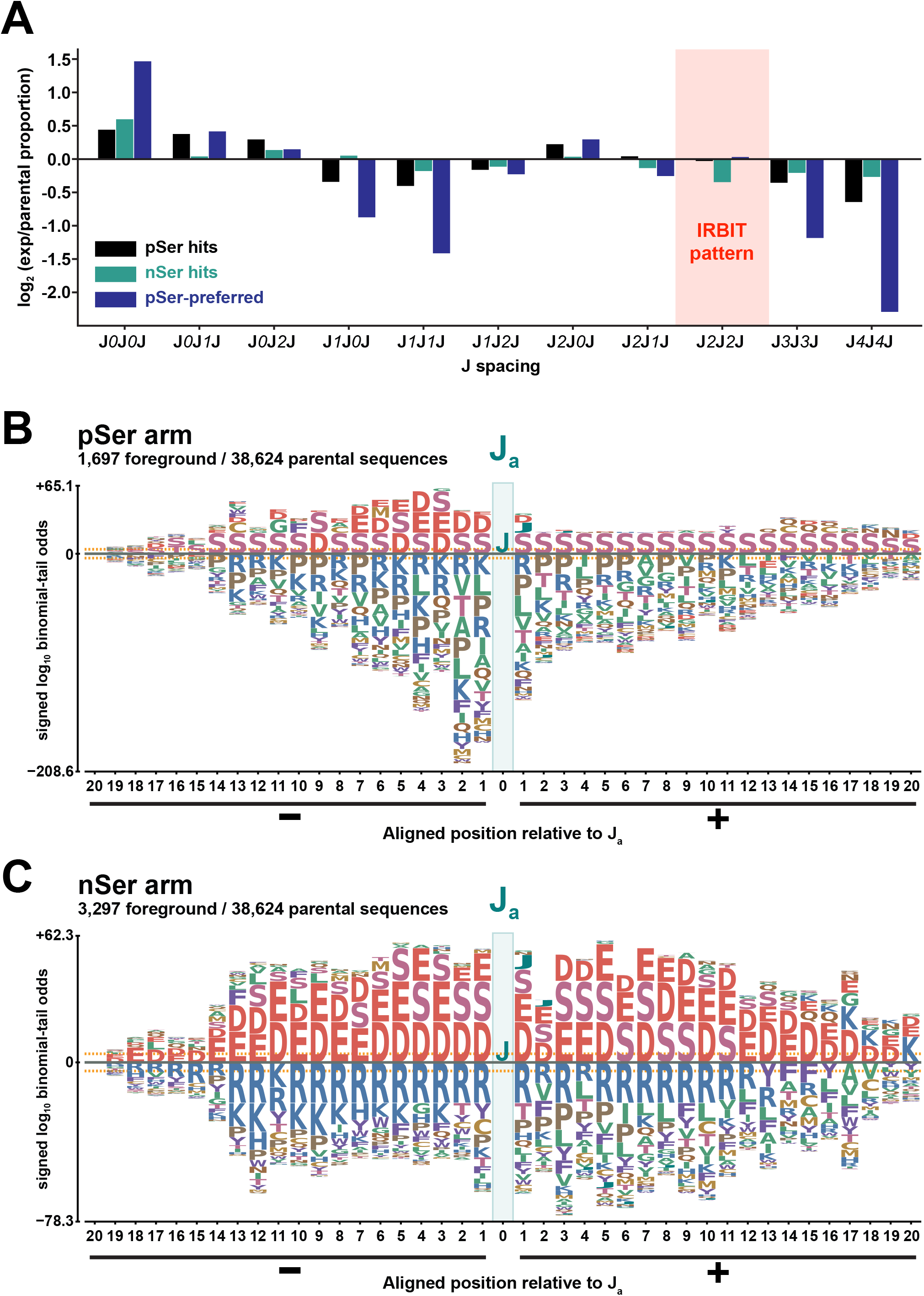
Phosphoserine spacing and local sequence context of peptides enriched in the PLCδ_1_-PH screen **A.** Relative representation of the 11 programmed three-site **J**-spacing classes among pSer hits (black), nSer hits (teal), and FDR-supported pSer-preferred sequences (blue). As described in the text, **J** denotes a site programmed to encode pSer or Ser. In **J***x***J***y***J**, *x* and *y* denote the numbers of residues between successive programmed UAG sites. Values plotted in the graph are log2 ratios of the proportion of each spacing-class in the different sets of hits compared to the proportion of entries belonging to that set in the designed parental library. Exact duplicate translated peptide sequences were removed independently from each set and background. Within-screen hits meet FC ≥2, q < 0.05, and nonzero selected-signal criteria. pSer-preferred entries meet the pSer within-screen hit criteria and additionally have a pSer/nSer enrichment ratio ≥2 and direct q < 0.05. The light red shaded region marks the IRBIT-associated **J***2***J***2***J** spacing that is not found to be enriched in any of the screens. (**B** and **C**) Probabilistic sequence logos^53^ for foregrounds in the pSer screen arm (**B**; 1,697 sequences) and nSer (**C**; 3,297 sequences) foregrounds, each compared with the 38,624-sequence designed parental background. These foregrounds require selected signal, FC ≥2, and unadjusted within-screen p < 0.05, followed by exact-sequence deduplication. Sequences are aligned at the first programmed site, **J**_a_, with positions −20 to +20 shown. Letters (amino acids) above or below the horizontal dotted zero line indicate overrepresentation or underrepresentation of that amino acid type, respectively. Letter heights reflect signed binomial-tail log_10_ odds. Dashed orange lines at approximately ±4.21 mark the Bonferroni threshold for familywise α = 0.05. Note that the panels use independent vertical scales.

Position-aware sequence analysis using pLogo^53^ further revealed substantial promiscuity in peptide binding to PLCδ_1_-PH. There were interesting differences between the array of peptide sequences that gave signal in the pSer screen (Figure 4B) and nSer screen (Figure 4C).

Sequences were aligned to the first UAG programmed site, **J**_a_, and exact duplicates of translated peptides were removed from each foreground and associated parental background. The pSer hits showed clear enrichment of aspartate/glutamate and depletion of lysine/arginine N-terminal to **J**_a_, particularly between positions-5 and-1, with no apparent charge requirement C-terminal to **J**_a_. nSer hits, by contrast, showed substantial selection for acidic residues across a broad sequence window – from-14 up to +14. These different characteristics are also evident when hits from the pSer and nSer screens are compared, with substantial enrichment of acidic residues from positions-5 to-1 before the aligned **J**_a_. Binding to PLCδ_1_-PH in Hi-P seems to depend on this positional pattern, rather than on charge itself, as pSer-preferred hits had essentially the same total D/E content as the parental sequences in the library (11.1% versus 10.7%). The charge distribution differences between pSer and nSer hits are also clear in Figure 5A and B. These data therefore suggest that PLCδ_1_-PH can recognize a particular pattern of negative charge – centered on a given phosphoserine (note that additional phosphoserines are not positionally conserved). When there is no phosphoserine, the peptides that bind most strongly appear to be those with the greatest negative charge. This analysis shows that the within-screen fold-enrichment for peptides increases substantially with mean D/E content across nSer hits, but only minimally across pSer hits.

**Figure 5.**
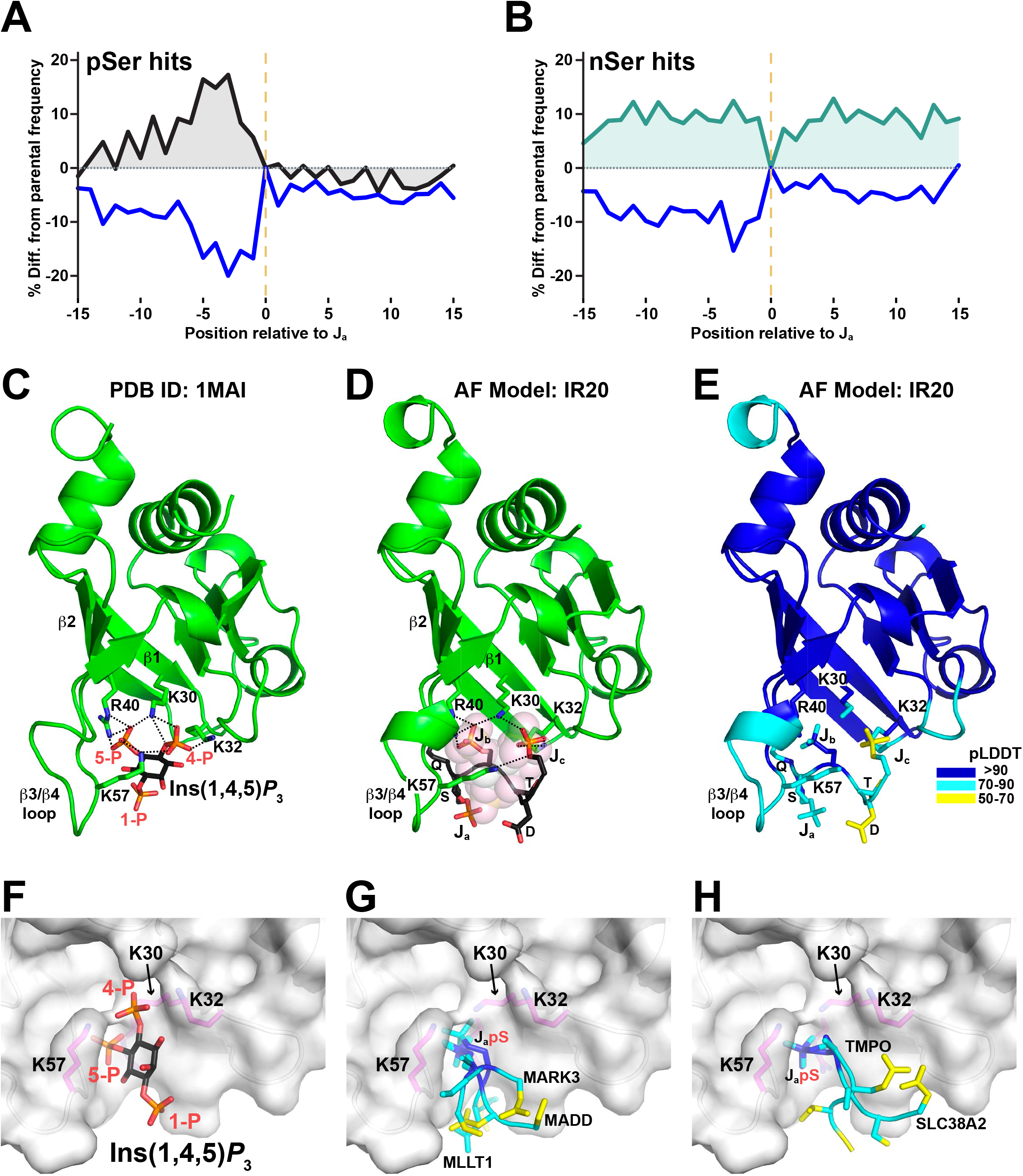
Promiscuity and specificity in peptide charge pattern recognition by PLCδ_1_-PH (**A** and **B**) Position-specific differences in combined acidic residue (D/E) and basic residue (K/R) frequencies between the pSer (**A**) or nSer (**B**) set of Hi-P hits compared with those in the designed parental library. Foregrounds are the q < 0.05 within-screen hit sets defined in Figure 3B and C, with exact duplicate translated peptides removed independently from each foreground and background. Sequences are aligned to **J**_a_, the first programmed UAG site. D/E curves are black in **A** and teal in **B**; K/R curves are blue. Values are percentage-point differences from parental frequencies. The vertical dashed line marks **J**_a_ at position zero. The horizontal dotted line marks no difference from the parental frequency. **C.** Crystal structure of PLCδ_1_-PH bound to Ins(1,4,5)*P*₃ (PDB: 1MAI), with the PH domain is shown as a green cartoon and the Ins(1,4,5)*P*_3_ ligand is shown as sticks, with oxygen atoms colored red and phosphorus orange. The ligand’s 1-, 4-, and 5-phosphate groups are labeled in red, and predicted polar contacts made by the 4-and 5-phosphates to the side chains of K30, K32, R40, and K57 are shown as dashed black lines. Side chains are shown as sticks, with nitrogen atoms colored blue. Strands β1, β2 as well as the β3/β4 loop are labeled. **D.** AlphaFold 3^55^ model of PLCδ_1_-PH bound to the triply phosphorylated (**J***2***J***2***J**) IRBIT-derived sequence SRSI**pS**QS**pS**TD**pS**, from the phosphoserine-rich N-terminal region. **J**_a_, **J**_b_, and **J**_c_ correspond to pS68, pS71 and pS74 of IRBIT. The predicted positions of the **J**_c_ and **J**_b_ phosphates correspond very closely to the positions of the 4-and 5-phosphates of Ins(1,4,5)*P*_3_ when bound, as shown with the transparent space-filling model of bound Ins(1,4,5)*P*_3_ in this figure. The position of **J**_a_ in the model is close to the crystallographically observed location of the Ins(1,4,5)*P*_3_ 1-phosphate. The IpTM and pTM scores^56^ for this AlphaFold3 prediction were 0.79 and 0.89 respectively, suggesting a high quality prediction – in which the three phosphates in the **J***2***J***2***J** pattern do appear to mimic the phosphate pattern in Ins(1,4,5)*P*_3_ – even though this is not a strongly selected sequence by Hi-P. **E.** The model from **D** is duplicated, with coloring dictated by predicted local confidence (pLDDT): dark blue, >90; cyan, 70-90; yellow, 50-70. **J**_b_, in the Ins(1,4,5)*P*_3_ 5-phosphate location, is positioned with the highest confidence. **F.** Surface representation of the binding pocket in PLCδ_1_-PH, occupied by Ins(1,4,5)*P*₃, in PDB ID: 1MAI.^12^ The protein surface is colored gray, with Ins(1,4,5)*P*₃ colored as in **C**, and K30, K32, R40 and K57 side chains shown as magenta sticks with blue nitrogens. Note that the Ins(1,4,5)*P*_3_ 5-phosphate penetrates a significant cavity in the PH domain surface, lined by basic side chains. **G** and **H**. Identical surface views of the PLCδ_1_ PH domain binding pocket as in **F**, but with top Hi-P singly phosphorylated peptides bound as modeled using AlphaFold 3. In **G**, the top hits described in the text from MARK3 (IpTM = 0.76), MLLT1 (IpTM = 0.83), and MADD1 (IpTM = 0.79) are shown. In each case, the **J**_a_ pSer phosphate group is predicted to lie in the PLCδ_1_-PH pocket that the Ins(1,4,5)*P*_3_ 5-phosphate occupies. The pLDDT value is at a minimum for the pSer side chain, suggesting confident prediction and specific recognition. Other residues, notably acidic side chains N-terminal to the **J**_a_ pSer vary in location, are less confidently predicted, and interacts with different basic side chains in a variety of orientations – suggesting delocalized attraction. **H.** Two other top Hi-P hits, from SLC38A2 (IpTM = 0.86) and TMPO (IpTM = 0.82) suggest a very similar arrangement. Peptides in G and H are colored by pLDDT as in **E**.

### Examples of top screen hits

Representative sequences enriched in one or both Hi-P screens provide some useful insight into the peptide recognition properties of PLCδ_1_-PH. The most recurrent pSer preferred region was from the MARK3/C-TAK1 kinase.^54^ Of 108 primary pSer preferred entries, 21 mapped to human MARK3 or the corresponding mouse protein, and all 21 of these represented distinct overlapping windows or programmed-site variants. One representative entry (**`TATYLLLGRKSSELDASDJJSSJNLSLAKVRPSSDLNNSTG`**) showed a 37.8-fold pSer enrichment (q=0.0057), but only a 1.1-fold nSer enrichment (q=0.872), consistent with the pLogo results in Figure 4B. An entry from super elongation complex subunit MLLT1 (**`MVEDLQSEESDEDDJJJGEEAAGKTNPGRDS`**) was strongly enriched in both screens, but favored pSer: pSer enrichment was 154.1-fold (q=0.0018) and nSer enrichment was also substantial at 17.2-fold (q=0.023). This shows four acidic side chains before **J**_a_, but two in the sequence that follows the **J***0***J***0***J** pattern. An entry from MAP kinase-activating death domain protein or MADD (**`SLRLASDSDAESDJRAJJPNSTVSNTSTEGF`**) showed strong pSer enrichment (38.2-fold, q=0.0012), but less nSer enrichment (1.43-fold, q=0.258), consistent with only having acid residues to the N-terminus of the **J***2***J***0***J** pattern. Unexpectedly, the 41-residue IRBIT-derived IR20 sequence that gave robust signals in our initial low-throughput Hi-P experiments with PLCδ_1_-PH (Figure 3A) – with sequence **`FTKFPTKTGRRSLSRSIJQSJTDJYSSAASYTDSSDDEVSP`** – was actually depleted in both screens (pSer FC 0.070, q=0.127; nSer FC 0.107, q=0.328), as was the nested 31-residue IR15 reporter with the same **J***2***J***2***J** IRBIT sequence. We do not consider that this result negates our results with IRBIT, but rather that it shows how a construct capable of producing fluorescence complementation when tested individually in Hi-P may fail to increase in relative abundance during pooled competitive selection. We also note that an entry from NOLC1, identified as a binding partner for the RASA1 PH domain with a **J***2***J***0***J** pattern (**`LSLPAKQAPQGSRDSSSDJDSJJSEEEEEKTSKSAVKKKPQ`**), showed some degree of nominal pSer enrichment (2.63-fold; p=0.0397, q=0.0870) although this result did not retain FDR support.

### Structural considerations

We used AlphaFold3^55^ to explore how PLCδ_1_-PH might recognize different phosphopeptides based on the Hi-P results with IRBIT IR20 and the broader screen. We first modeled PLCδ_1_-PH bound to the **J***2***J***2***J**-containing sequence **`SRSIpSQSpSTDpS`**, which yielded a complex with an interface predicted template modeling (ipTM) score^55,56^ of 0.8 – suggesting a confident high-quality prediction. For comparison, an AlphaFold3 prediction of the PLCδ_1_-PH/Ins(1,4,5)*P*_3_ complex has an ipTM score of 0.93. Figure 5C-E shows how **J**_b_ and **J**_c_ of the multiply phosphorylated IRBIT peptide are predicted to occupy close to the same positions as the 5-and 4-phosphates of Ins(1,4,5)*P*_3_ in the crystal structure of the PLCδ_1_-PH/Ins(1,4,5)*P*_3_ complex.^12^ Key basic side chains in the **KX_n_[K/R]XR** motif (K30, K32, R40 in the β1/β2 loop), plus K57, are engaged by the modeled IR20 peptide (Figure 5D) just as seen for Ins(1,4,5)*P*_3_ (Figure 5C). The **J**_a_ pSer is also close to where the 1-phosphate of Ins(1,4,5)*P*_3_ is located. This arrangement thus reveals how it is possible in principle that a **J***2***J***2***J** pattern as seen in IRBIT may place the three phosphate groups in locations closely resembling those presented in Ins(1,4,5)*P*_3_ – although this was far from the preferred peptide in Hi-P.

As described above, contrary to our initial expectations, we did not see conserved patterns of multiple pSer/**J** locations by Hi-P. Our initial experiments with IRBIT sequences did suggest that multiple pSers were required for binding of that sequence, which lacks acidic residues N-terminal to the **J***2***J***2***J** pattern. By contrast, the pLogo output in Figure 4B suggests that just one pSer is sufficient, as long as it is preceded by several residues with acidic side chains. AlphaFold modeling of the top peptides that meet this criterion mentioned in the previous section – from MARK3, MLLT1, MADD – as well as two other frequent pSer hits from SLC38A2 and thymopoietin is intriguing. In each case, AlphaFold placed the bound peptide in the crystallographically defined Ins(1,4,5)*P*_3_ binding pocket of PLCδ_1_-PH,^12^ with the **J**_a_ phosphate buried in a pocket occupied by the 5-phosphate in the Ins(1,4,5)*P*_3_ complex. This pocket is bounded by the side chains of K30, R40 and K57 in the PLCδ_1_ PH domain. The ipTM scores for these models ranged from 0.76 (for the MARK3 entry) to 0.86 (from the SLC38A2 entry). These peptides – and presumably most of the peptides responsible for signal in the pSer arm of our Hi-P experiments – are likely to gain substantial binding energy by placing the key phosphate group of their **J**_a_ pSer in this somewhat buried pocket. The acidic side chains that precede the **J**_a_ pSer interact with other basic and hydrophilic side chains in the PLCδ_1_-PH binding site, including K32, which is not engaged by the buried pSer side chain, R38, and R46 that are reoriented in the AlphaFold models to accommodate the peptides. Comparison of these complexes suggests that peptide recognition occurs through a combination of pSer accommodation in a well-defined basic pocket in the PH domain plus delocalized (and less stereospecific) electrostatic attraction in other regions of the interface. This view is consistent with the finding that peptides preferred in the nSer arm of the Hi-P screen gave signal if they were acidic enough. Our data suggest that PH domains like that from PLCδ_1_ have a propensity to associate with negatively charged regions in proteins (Figure 4C), with distributed acidic residues supporting a weak encounter. Phosphorylation appears to increase local charge density, extend anionic contacts, and/or favor a subset of presentations. Interestingly, this finding is highly reminiscent of results obtained with the Tfb1/p62 PH-like domain,^57^ which binds an acidic transactivation domain (TAD) region of p53 with the sequence **`SPDDIEQWFTE`** in a way that is greatly enhanced by phosphorylation of the serine and threonine at the beginning and end of the sequence.^28^ These peptides bind to a positively charged region on the surface of the p62 PH-like domain, although it is in a different location on the β-sandwich than the basic surface of PLCδ_1_-PH. Another close analogy can be drawn with the CERT PH domain, which shares homology with the phosphoinositide-binding FAPP1^31^ and Osh1/2p^31^ PH domains. In its active state, CERT associates through its PH domain with PtdIns(4)*P* in the Golgi membrane. At that membrane, CERT becomes hyperphosphorylated in a region adjacent to the PH domain^58^ that associates intramolecularly with the CERT PH domain to displace it from the Golgi membrane and inhibit ceramide transfer activity.^29^

## CONCLUSIONS

Our findings provide a clear structural rationale for why acidic stretches of proteins have frequently been reported as ligands for PH and PH-like domains.^17,25,27,29,58^ Although we initially considered that these might represent alternative ligands for PH domains that do not bind to phosphoinositides, we found in *S. cerevisiae* that more PH domains bind the acidic Nsr1p nucleolin equivalent, and that all phosphoinositide-binding yeast PH domains^18^ also bind Nsr1p. This suggests that phosphoinositides or their soluble inositol phosphate moieties bind similarly to the same sites. Indeed, a highly phosphorylated stretch of IRBIT was similarly shown to compete with Ins(1,4,5)*P*_3_ for binding to the structurally unrelated Ins(1,4,5)*P*_3_ receptor.^36,37,46^ Our binding studies *in vitro* and in bacteria argue that a multiply phosphorylated region within the IRBIT N-terminus requires multiple phosphorylations to bind PLCδ_1_-PH, just as described for its Ins(1,4,5)*P*_3_R binding^36^ – and centered on the same phosphoserines (S68, S71, and S74). In this setting, the **J***2***J***2***J** sequence pattern appears to mimic Ins(1,4,5)*P*_3_ stereochemically, placing its three phosphate groups in very similar locations. Although this interaction was confirmed using the Hi-P platform in a low throughput manner – and its phosphorylation dependence was clear – the multiply phosphorylated IRBIT peptide was not highly selected for in our Hi-P screen. Rather, what dominated in this context were peptides with a single pSer, preceded by concentrated negative charge. Structural analysis of how these peptides might bind using AlphaFold suggests that the single phosphate group of the peptide projects into the partly buried pocket in the PH domain that plays a crucial role in accommodating the 5-phosphate in the PLCδ1-PH/Ins(1,4,5)*P*_3_ complex and the 3-phosphate in complexes of 3-phosphorylated inositol phosphates with PH domains that bind PI 3-kinase products.^10,11,14^ The remaining acidic regions of the bound peptide engage in more delocalized interaction with basic side chains at the binding pocket that have substantial flexibility and can accommodate different negatively charged ligands. Thus, peptide recognition involves a combination of specific phosphate recognition (in the 5-P pocket shown in Figure 5F-H) and delocalized association with acidic stretches of amino acids. Even in the absence of phosphorylation, our Hi-P data suggest substantial binding to regions that are simply rich in glutamates and aspartates.

Phosphorylation may enhance or specify PH domain interaction with such regions, as suggested by studies with p53 binding to the p62 PH-like domain in TFIIH^28^ and with CERT – for which the PH domain binds PtdIns(4)*P* or an adjacent phosphorylated part of the same protein.^29,58^ Along the same lines, recent work with IRBIT binding to the Ins(1,4,5)P_3_R^37^ has suggested that more than one phosphorylated region may be able independently to compete with (and mimic) Ins(1,4.5)*P*_3_ binding to the receptor. This may reflect a similar promiscuity in binding of a multiply serine-phosphorylated protein region to Ins(1,4,5)P3 as we see for peptides binding to PLCδ_1_-PH.

All of the PH domains that we have studied here bind phosphoinositides with quite high affinity, and acidic peptides seem to represent alternative ligands – subject to the criteria uncovered in our screen. However, this class of PH domains represents only a small minority, with most PH domains – at least in *S. cerevisiae* – failing to bind strongly to phosphoinositides.^18^ Nearly all PH domains have substantial electrostatic sidedness,^18,20^ with classes that differ in the location of the most basic region. One example is the spectrin PH domain, which binds Ins(1,4,5)*P*_3_ at a site on the opposite side of the β1/β2 loops from that seen for PLCδ_1_-PH. We suggest that other PH domains, with basic regions in different locations and with different structural characteristic, may bind acidic and/or phosphorylated regions of proteins with different features. Extending the work described here for PLCδ1-PH across other PH domain classes may thus provide new information on the functions of poorly understood examples of this common domain class.

## RESOURCE AVAILABILITY

### Lead contact

Requests for further information or reagents may be directed to the lead contact, Mark A. Lemmon.

### Materials Availability

All unique and stable reagents generated in this study are available upon request from the lead contact.

### Data and Code Availability

- All data reported in this paper will be shared by the lead contact upon request.
- Raw FCS and FASTQ files, processed entry-level analysis tables, and source data underlying the displayed results are held by the authors and are available from the lead author upon reasonable request.
- PDB entry 1MAI was used to generate images in Figure 1A, Figure 5C, and Figures 5F-H.
- AlphaFold was used to generate structural predictions in Figure 5.
- This paper does not report original code.
- Any additional information required to analyze or reanalyze the data reported in this paper is available from the lead contact upon request.

## ACKNOWLEDGMENTS

We thank the members of the Lemmon and Ferguson labs for thoughtful discussions, the Rinehart laboratory at the Yale Systems Biology Institute for sharing the Hi-P platform, and Maya Kornaj for assistance with initial platform setup and training. We also thank the Yale Center for Genome Analysis for next-generation sequencing services. This work was supported by NIH grant R35-GM122485 (to M.A.L.).

## AUTHOR CONTRIBUTIONS

Conceptualization, C.K.N. and M.A.L.; Methodology, C.K.N., J.M.M., J.W.M.; Investigation, C.K.N., J.M.M., J.W.M., M.S.; Resources, S.E.S., M.A.L., K.M.F.; Visualization, C.K.N. and M.A.L.; Writing – Original Draft, C.K.N. and M.A.L.; Writing – Review and Editing, C.K.N., M.A.L., K.M.F., S.E.S., J.M.M., J.W.M., and M.S.; Funding Acquisition, M.A.L.; Supervision, M.A.L. and K.M.F.

## STAR Methods

Detailed methods are provided in the online version of this paper and include the following:

- KEY RESOURCES TABLE
- **EXPERIMENTAL MODEL AND STUDY PARTICIPANT DETAILS**

`o` Cell lines and culture conditions
- METHODS DETAILS

`o` IRBIT and PH-domain constructs
`o` GST and GST/PH expression and purification
`o` Yeast PH domain expression and pull-down experiments
`o` Sf9 cell expression and purification of full-length IRBIT
`o` Surface-plasmon resonance (SPR) binding studies
`o` Biolayer Interferometry (BLI) concentration-series measurements
`o` BLI soluble-ligand perturbation measurements
`o` Hi-P reporter architecture and low-throughput measurements
`o` Triple-site library design and analytical units
`o` Library cloning, transformation, and representation
`o` Measurement of mCherry fluorescence complementation and cell sorting
- QUANTIFICATION AND STATISTICAL ANALYSIS

`o` Amplicon sequencing and read mapping
`o` Normalization, statistical testing, and hit calling
`o` Sequence-feature and charge-context analyses
`o` Analysis software

## STAR Methods

### EXPERIMENTAL MODEL AND STUDY PARTICIPANT DETAILS

#### Cell lines and culture conditions

*Spodoptera frugiperda* Sf9 cells (Expression Systems), originally established from immature ovaries of female *S. frugiperda* pupae, were propagated at 27°C with constant shaking at 120 rpm in either serum-free ESF 921 Insect Cell Culture Medium (Expression Systems) containing 50 U/mL penicillin/streptomycin or Insect-XPRESS medium (Lonza), and were used for production of all proteins. *E. coli* BL21(DE3) cultures were grown in 1 L Luria Broth (LB) containing 150 μg/mL ampicillin at 37°C for protein production. C321.ΔA *E. coli* cultures for Hi-P screens were grown in LB at 25°C with constant shaking. TAP-tagged proteins were expressed in the S288C strain of *S. cerevisiae* (MATa his3Δ1 leu2Δ0 met15Δ0 ura3Δ0) used for constructing the yeast TAP-tagged open reading frame (ORF) collection.^39^

## METHODS DETAILS

### Methods

#### IRBIT and PH-domain constructs

Human IRBIT (AHCYL1; UniProt reference O43865) was studied as the full length 530 residue protein expressed with an N-terminal hexahistidine tag. The corresponding DNA was PCR amplified and subcloned into the Bam HI and Xho I restriction sites of pFastbacHT A (Invitrogen/Life Technologies) for expression using the Bac-to-Bac system in Sf9 cells. Focused mutated variants were generated by site-directed mutagenesis using QuikChange, and verified by Sanger sequencing. The focused M1-M5 variants included M1 (S68A/S71A/S74A), M2 (S71A/S74A/S77A), M3 (S66A/S68A/S70A), M4 (S70A/S74A/S77A), M5 (S66A/S70A/S77A), IRBIT-A_7_ (S66A/S68A/S70A/S71A/S74A/S76A/S77A) and IRBIT-E_7_ (S66E/S68E/S70E/S71E/S74E/S76E/S77E). The glutathione-S-transferase (GST) fused PLCδ_1_ PH domain (GST/PLCδ_1_-PH) comprised rat PLCδ_1_ residues 11-140 in the pGEX-2TK construct reported by Kavran *et al*.^42^ and was expressed as described. The GST-fused human dynamin-1 (Dyn1) PH domain was expressed as described by Klein *et al*.^44^ and served as a low-affinity negative control for PH domain phosphoinositide binding studies. *S. cerevisiae* PH domains were expressed as GST fusion proteins using the constructs described by Yu *et al*.^18^ The yeast TAP-fusion collection^39^ and TAP tag antibody were purchased from Thermo Scientific/Open Biosystems. Glutathione agarose beads and GST antibody were purchased from Sigma. His5-binding antibody was obtained from Qiagen. The Hi-P domain-side reporter placed GST/PLCδ_1_-PH N-terminal to mCherryC as for other phosphopeptide binding domains in previous studies.^47,48^

#### GST and GST/PH expression and purification

*E. coli* BL21(DE3) cultures expressing GST or GST/PH fusions were grown in 1 L LB containing 150 μg/mL ampicillin at 37°C to OD_600_ 0.6-0.8, induced with 1 mM IPTG, and incubated for a further 2 h at 37°C (or 25°C where specified by Yu *et al*.^18^ for yeast PH domains). Pellets were thawed on ice and lysed in 50 mM Tris, pH 8.0, containing 150 mM NaCl, 5 mM DTT, 1 mM EDTA, 1 mM PMSF, and 0.5 mg/mL lysozyme. Samples were sonicated on ice for 5 min total at 30% power using 2 s-on/3 s-off pulses and clarified by centrifugation at >90,000 x g for 75 min at 4°C. Clarified material was batch-bound to Pierce glutathione agarose (approximately 200 μL settled beads/400 μL slurry per preparation) for 1-1.5 h at 4°C with mixing. Resin was washed five times with approximately 1 mL 50 mM Tris, pH 8.0, 150 mM NaCl, and 1 mM EDTA. Protein was then eluted in three fractions of approximately 500 μL each in the same buffer containing 10 mM reduced glutathione. Proteins concentration was measured by A_280_ after tenfold dilution. Proteins were retained at 4 C in this buffer and used within 1 week. Separate physical preparations were used across experiments under the same protocol.

#### Yeast PH domain expression and pull-down experiments

*S. cerevisiae* cells expressing the relevant TAP-tagged proteins were harvested as described^59^ and lysed by bead beater according to the manufacturer’s instructions (Biospec Products). Briefly yeast cells were resuspended in 50 mM Tris-HCl, pH 8.0, containing 150 mM NaCl, 1 mM PMSF, 1 mM DTT, 10 μM benzamidine, 2.3 μM leupeptin, 2 μM aprotinin, 3 μM pepstatin, 30 mM NaF and 100 mM Na_2_MoO_4_ and placed in a prechilled 50 mL lysis chamber containing 0.5 mm glass beads filled to above the rotor blades (∼1/3 full). The cell suspension was pulsed for 7 cycles of 30 s each and filtered through coarse Whatman paper to remove the beads. Lysates were then clarified by centrifugation at 17,000 rpm for 20 min.

Soluble yeast lysate expressing TAP-tagged *S. cerevisiae* Nsr1p was divided into equal parts and incubated with normalized levels of GST/PH fusion protein bound glutathione sepharose beads. The lysates were incubated with GST/PH fusion for 2 h at 4°C, after which the beads were washed five times with the yeast lysis buffer, treated with denaturing protein sample buffer and boiled prior to running on SDS-PAGE gels. Western blots were probed with the relevant antibodies according to manufacturer recommendations.

#### Sf9 expression and purification of full-length IRBIT

Sf9 suspension cultures in ESF 921 were infected at approximately 2 x 10^6^ cells/mL with 25 mL P2 baculovirus suspension per liter and incubated at 27°C with shaking at 120 rpm. Cells were maintained below passage 15. Viral titer was not measured; the inoculum is therefore not expressed as a multiplicity of infection. Cultures were harvested at approximately 90% viability, collected by centrifugation at 3,000 x g, washed with phosphate-buffered saline (PBS), and frozen. Pellets were lysed in 20 mM HEPES, pH 7.5, containing 150 mM NaCl, 5 mM imidazole, 5 mM β-mercaptoethanol, 10% (w/v) glycerol, and 1 mM PMSF, without detergent or phosphatase inhibitor. Samples received approximately 3 kJ sonication on ice and were clarified by centrifugation at >90,000 x g for 75 min at 4°C. Clarified material was batch-bound to approximately 1-1.5 mL Qiagen Ni-NTA agarose for 45 min at 4°C. The resin was then washed with 22.5 mL of the same lysis buffer containing 5 mM imidazole. Subsequent elutions contained 20, 200, and 400 mM imidazole, respectively at pH 7.5; two 2 mL 200 mM imidazole fractions were collected for use in experiments, treated with 1× cOmplete EDTA-free protease inhibitor, and kept at 4°C for use within 1 week. Before binding studies, IRBIT proteins were desalted in 2 mL Zeba columns (approximately 650 μL loaded) and GST fusion proteins were desalted in 0.5 mL Zeba columns (approximately 130 μL loaded) into 20 mM HEPES, pH 7.5, containing 150 mM NaCl, 3 mM EDTA, and 0.005% Tween-20 (HBS-EP). Proteins were passed through Costar Spin-X 0.22 μm cellulose-acetate filters prior to use, and protein concentration measured by A_280_.

#### Surface-plasmon resonance (SPR) binding studies

SPR binding experiments were performed using a BIAcore 3000 instrument at 25°C using a running buffer of 10 mM HEPES buffer, pH 7.4 that contained 150 mM NaCl, 3.4 mM EDTA and 0.005% Tween-20. The noted His_6_-tagged IRBIT proteins were purified and coupled to BIAcore CM5 sensor chips by standard amine coupling (GE Healthcare). Briefly, the hydrogel matrix of BIAcore CM5 Biosensor chips was activated with N-hydroxysuccinimide (NHS) and *N*-ethyl-*N’*-[3-(diethylamino)propyl] carbodiimide (EDC). IRBIT in 10 mM sodium acetate, pH 4.8 was then flowed over the activated surface at 5 mL/min for 10 min. Non-crosslinked IRBIT was then washed away, and unreacted sites were blocked with 1 M ethanolamine, pH8.5. Increasing concentrations of purified wild-type GST/PLCδ_1_-PH were then flowed over the resulting surfaces. GST-PH fusion concentration was then plotted against percent saturation of the IRBIT surface, calculated by subtracting the SPR response on the control mock-coupled surface from the SPR response on the experimental surface and normalizing to the calculated B_max_ using one-site binding analysis in GraphPad Prism software.

#### Biolayer interferometry (BLI) concentration-series measurements

All displayed BLI experiments were performed using a PALL FortéBio Octet RED96e BLI system, employing Octet Anti-GST Biosensors. GST, GST/PLCδ1-PH, and GST/Dyn1-PH were loaded on to sensors. Anti-GST biosensors (Sartorius, catalog no. 18-5096), supplied precoated with GST antibody, were hydrated in HBS-EP according to the manufacturer’s instructions.

GST-tagged PH domains or GST alone were captured by immersing the sensors in 200 nM protein solutions in HBS-EP for 5 min. Loaded GST/PLCδ_1_-PH sensors were transferred into soluble IRBIT-WT, IRBIT-A_7_, or M1-M5 at a range of concentrations from 0.23 μM to 7.5 μM (depending on experiment) in HBS-EP buffer, in steps of approximately 2-fold for the concentration series, with a well volume of 150 μL. Corresponding traces for sensors with GST-only immobilized were subtracted, and GST/Dyn1 PH served as a low-binding comparator. The BLI response at each non-zero IRBIT protein concentration was taken as the mean over the final 20 s of association. Each experiment was fit independently by unweighted least squares to the equation: y (response) = B_max_[x/(*K*_d app_ + x)], without parameter constraints or outlier removal. For cross-experiment visualization, responses were divided by the within run mean fitted B_max_ value obtained with the strongest binders (WT and M5 IRBIT). This vertical scaling did not alter *K*_d app_. Profile-likelihood 95% confidence intervals were calculated for the run level fits.

#### BLI soluble-ligand perturbation measurements

Anti-GST sensors loaded with GST/PLCδ_1_-PH or GST alone were measured in matched cycles containing soluble Ins(1,4,5)*P*_3_ alone and ligand plus full-length wild-type IRBIT. Traces were aligned at the onset of association. The matched GST-only trace was subtracted to remove nonspecific IRBIT association with GST, and the corresponding ligand-only response was then subtracted. Endpoints were final 20 s means. Ins(1,4,5)*P*_3_ was tested at 25 μM with 5 μM and 1 μM wild-type IRBIT in three independent experiments. One sensor represented each within-run condition. These measurements quantify reduced GST-subtracted association under the tested geometry and do not determine an inhibition constant or common binding site.

#### Hi-P reporter architecture and low-throughput measurements

Hi-P experiments were implemented in C321.ΔA cells as an adaptation of experiments described by Barber *et al*.^47^ The pooled-screen reporter orientation involved co-expression of mCherryN-peptide paired with GST/PLCδ_1_-PH-mCherryC from a bicistronic plasmid. The same reporter-plasmid library was introduced separately into cells carrying SepOTSλ, to encode pSer at the three UAG sites, or the *supD* serine-suppressor tRNA plasmid, to encode serine (nSer) at those positions. The pSer and nSer screen arms were independent biological series. Low throughput reporter cultures were evaluated under uninduced, L-arabinose-only, and L-arabinose-plus-anhydrotetracycline (ATC) induced conditions. The selected pooled-screen condition was 0.067% L-arabinose plus 900 ng/mL ATC followed by overnight incubation with shaking at approximately 25°C. For the displayed benchmark experiment, mean fluorescence from LB-only wells was subtracted and values were divided by 1,000. Bars show means and SD across nine technical wells per condition. Technical wells were not treated as biological replicates.

#### Triple-site library design and analytical units

The 3pSerOligoLibraryDesign workflow mined documented human, mouse, and rat phosphoserine sites from PhosphoSitePlus^52^. Sites were filtered into 11 prespecified three-position classes: the nine **J***x***J***y***J** combinations for *x* and *y* equal to 0, 1, or 2 intervening residues, plus **J***3***J***3***J** and **J***4***J***4***J**. Approximately ±15-and ±20-residue windows were generated when sequence context permitted. Each design was codon optimized for bacterial expression, the three target serines were replaced with UAG codons, and cloning adapters were added. The final annotation table contained 41,848 source-specific entries. The synthesis record contained 38,624 unique 109-184-bp oligonucleotide designs, each ordered in duplicate (Agilent). Exact translated peptide sequences were used for independent deduplication in sequence-level summaries, and case-normalized gene labels were used only for gene-count summaries and clustered standard errors. The annotation entry remained the statistical unit for normalized abundance, replicate enrichment, and hit calling.

#### Library cloning, transformation, and representation

The pooled oligonucleotide library was ligated into the bicistronic Hi-P vector and introduced into Electromax DH10B cells for high-efficiency library construction. A representative 10^5^-fold dilution plate contained 343 colonies, corresponding to an estimated 3.43 x 10^7^ vector-library transformants. This equals population-level averages of approximately 820 transformants per annotation entry and 890 per unique oligonucleotide design; note that these averages do not imply uniform abundance. Plasmid recovered from the pooled DH10B transformation was introduced into C321.ΔA cells carrying SepOTSλ or *supD*. Every recovered expression library population contained at least 10^7^ colony forming units (CFU). Four unsorted transformed cell samples contained non-zero normalized sequence abundance values for 40,698, 40,689, 40,612, and 40,771 of 41,848 annotated entries. Each selected sample was compared with its matched unsorted sibling to account for non-uniform starting abundance.

#### Measurement of mCherry fluorescence complementation and cell sorting

For low throughput Hi-P assays, individual reporter cultures were induced with 0.067% L-arabinose and 900 ng/mL ATC and incubated overnight with shaking at 25°C, unless otherwise indicated. mCherry fluorescence was measured directly for these experiments in growth medium using 200 μL aliquots in black, flat-bottom 96-well plates using a BioTek Synergy 2 microplate reader, with excitation and emission filters of 585/10 nm and 620/15 nm, respectively (center wavelength/bandwidth). Mean fluorescence from LB-only wells was subtracted from each sample. For OD-normalized measurements, blank-corrected fluorescence was divided by the corresponding OD_620_ after subtraction of the mean LB-only OD_620_ value. mCherry was measured at the single cell level for Hi-P screens using fluorescence-activated cell sorting (FACS). Data acquisition and FACS sorting were performed using a BD FACSAria III SORP using BD FACSDiva 9.5.1. mCherry was recorded as PE-CF594-A at 650 V. All 24 screen files used identical detector voltages and an FSC acquisition threshold of 5,000. The gate sequence applied FSC/SSC bacterial and debris exclusion, pulse-geometry doublet discrimination, and the mCherry selection gate. For each selection round, the top 2% of mCherry events were collected with a target of 100,000 events. Sorted cells were expanded, re-induced, and subjected to a second top 2% collection. The doubly selected population was then expanded for plasmid isolation and insert sequencing. A small re-induced aliquot was analyzed by FACS after round two to document the final population; it was not subjected to selection and did not constitute a third round.

The four independent pSer screens were designated E1-E4 and the four independent nSer screens D1-D4. Each transformation yielded an unsorted sibling retained as its matched parent. Pairing was within screen arm and experiment: selected E1 was compared with parental E1, selected E2 with parental E2, and so forth. E1-E4 and D1-D4 were independent biological series.

## QUANTIFICATION AND STATISTICAL ANALYSIS

### Amplicon sequencing and read mapping

Plasmid DNA was isolated from each doubly selected Hi-P population and its matched unsorted sibling. Peptide-library inserts were amplified by PCR and submitted to the Yale Center for Genome Analysis for next generation sequencing. The four barcoding-primer pairs were MAK084/MAK085, MAK086/MAK087, MAK088/MAK089, and MAK111/MAK112. Sequencing used NovaSeq paired-end 150-bp reads with dual i5/i7 indexing, and paired R1/R2 FASTQ files were delivered. FASTQ files were processed against seq_list_ref.fasta, which contained 41,848 reference entries. The workflow generated BBMap PileupStats.txt coverage output^60^ and samtools idxstats alignment output.^61^ Sequence abundance was taken from the PileupStats Avg_fold field, which reports average fold coverage.

### Normalization, statistical testing, and hit calling

Within each sample, Avg_fold values were divided by their sum across all library entries and multiplied by 1,000,000. These quantities are normalized sequence-abundance values, not raw read counts. For a matched selected-parental replicate *i*, enrichment was calculated as d*_i_*= log2[(selected*i* + 0.5)/(parental*i* + 0.5)]. The pseudocount of 0.5 normalized abundance units was applied symmetrically to keep the ratio finite when one value was zero. Informative-pair eligibility was established before pseudocount application. A pair was informative when both measurements were available and at least one side was nonzero; pairs with zero on both sides were excluded. At least three informative pairs were required for inference. The displayed within-screen fold enrichment was FC = 2^(mean d*_i_*), the geometric mean of replicate-level fold enrichments. Within each screen arm, a two-sided one-sample t test of the replicate-level log2 enrichment values against zero was applied to entries with at least three informative pairs and nonzero variance. All eligible finite *p* values entered the arm-specific Benjamini-Hochberg family before the selected-signal gate. A within-screen hit required FC≥2, q<0.05, and a nonzero normalized sequence-abundance value in at least one selected replicate. The direct comparison used a two-sided Welch t test on the pSer and nSer replicate-enrichment vectors because the screen arms were independent. Direct testing required at least three informative pairs in each arm. Benjamini-Hochberg correction was applied across all eligible direct p values. The displayed direct pSer-versus-nSer ratio was FCpSer/FCnSer. An FDR-supported pSer-preferred entry was required to be a pSer within-screen hit, have a direct pSer-versus-nSer ratio of at least 2, and have direct q<0.05. An all-zero selected condition was not treated as observed enrichment. It could retain finite p and q values when the informative-pair, variance, and finite-test criteria were satisfied because it represented a testable depletion. For display, an all-zero selected condition was assigned FC=0. Displayed direct ratios of 0 or infinity were descriptive and not interpreted as measured zero or infinite effects.

### Sequence-feature and charge-context analyses

Exact translated-peptide duplicates were removed independently from each foreground and its applicable parental background for sequence-level analyses. Spacing representation was calculated as the proportion of each **J**-spacing class in a selected set divided by its proportion in the deduplicated designed parent and displayed on a log2 scale. All 11 classes were retained. Position-aware residue analyses aligned sequences at the first programmed site, **J**_a_, across positions-20 to +20. Terminal positions with incomplete coverage used position-specific denominators. Differences in combined D/E or K/R frequency were calculated relative to the deduplicated designed parental library. The motif-oriented supplementary analysis used Active-

Gate, a custom renderer implementing the pLogo statistic relative to the designed parental background^53^. Its pSer-preferential foreground contained 113 exact peptide sequences defined by pSer FC≥2, raw pSer p<0.05, direct ratio≥2, and direct q<0.05. This visualization set is distinct from the primary 108-entry FDR-supported pSer-preferred set. For each annotation entry, global acidic content was the fraction of encoded residues that were D or E. At within-screen FC thresholds of at least 2, 4, 8, 12, 16, and 20, entries were retained only when q<0.05 and selected signal was present. The figure reports entry-weighted means, SEM, and retained entry counts. The threshold-defined sets are nested. At each threshold, D/E fraction was regressed on screen arm with standard errors clustered by case-normalized gene label; the six two-sided P values were Bonferroni adjusted. Among FC≥2 hits, a linear model containing log2FC, arm, and their interaction tested whether the acidic-content relationship differed between screens, again using gene-clustered standard errors.

### Analysis software

Entry-level annotations, processed abundances, enrichments, and statistical outputs were generated using the analysis workflow described above. Sequence-feature analyses and the interactive screen explorer used the same analysis outputs.

